# Mild Hyperthermia Enhanced Liposomal Doxorubicin Delivery and CD8^+^ T cell Infiltration in Triple Negative Breast Cancer

**DOI:** 10.1101/2024.04.25.591226

**Authors:** Farzaneh Rezazadeh, Wajfa Saadat, Ryan Smith, Alexander Pattyn, Mohammad Malik, Fuad Yazdani, Allen-Dexter Saliganan, Mohammad Mehrmohammadi, Nerissa T. Viola

## Abstract

Mild hyperthermia (MHTh) is often used in combination with chemotherapy and radiotherapy for cancer treatment. In the current study, the effect of MHTh on the enhanced uptake of the FDA-approved chemotherapy drug, liposomal doxorubicin (dox) in syngeneic 4T1 tumors was investigated. Doxorubicin has inherent fluorescence properties having an emission signal at 590 nm upon excitation with a 480 nm laser. A group of mice administered with doxorubicin (dox) were exposed to MHTh (42 °C) for 30 minutes whereas control group given dox did not receive MHTh. *Ex vivo* optical imaging of harvested tumors confirmed higher uptake of dox in treated versus the control untreated tumors. Confocal microscopy of tumor sections indicates higher fluorescent intensity due to increased accumulation of dox in MHTh-treated compared to untreated tumors. We examined the effect of MHTh to enhance CD8 tumor infiltration, production of interferon-γ (IFN-γ) and expression of programmed death ligand-1 (PD-L1). mRNA *in situ* hybridization was performed to test for transcripts of CD8, IFN-γ and PD-L1. Results showed that higher expression of CD8 mRNA was observed in MHTh-administered tumors versus untreated cohorts. The signal for IFN-γ and PD-L1 in both groups were not significantly different. Taken together, our findings imply that MHTh can improve tumor uptake of dox. Importantly, our data suggests that MHTh can boost CD8^+^ T cell infiltration.

## Introduction

Breast cancer is the second most common leading cause of death in women [1]. The most aggressive type of breast cancer is triple negative breast cancer (TNBC) because it lacks the expression of commonly targeted receptors, estrogen, progesterone, and human epidermal growth factor receptor 2 (HER-2) [2, 3]. Three standard therapeutic methods for the treatment of breast cancer include surgery, chemotherapy, and radiotherapy [4, 5]. The most effective chemotherapeutic drug used to treat TNBC is doxorubicin [6]. However, a major challenge of the conventional chemotherapy is related with considerable side-effects [7]. Many attempts have been made to overcome this challenge by changing the biodistribution of drugs in favor of tumor accumulation and reduce toxicity. Encapsulating doxorubicin within liposomal nanoparticle is one of the solutions to prolong drug circulation and reduce side effects [8, 9]. However, the efficacy of the drug was not substantially improved due to slow and passive drug release from the liposomes [10, 11].

Mild hyperthermia (MHTh) is a therapeutic procedure by which a tumor is locally heated to rises to a higher level than normal (∼42 °C) has been developed as an anticancer treatment for several decades [12-14]. Hyperthermia can directly cause cytotoxic effect mainly based on denaturation of nucleotide, cytoplasmatic or membrane proteins or by sensitizing the cells to other treatment modalities like radiation and chemotherapy [15]. MHTh is routinely used in the clinic as an adjuvant therapy and is known to increase the effectiveness and greatly reduce the dosage requirement of chemotherapy agent like doxorubicin [13, 16, 17]. Mechanisms by which MHTh can improve the efficacy of chemotherapy include increased perfusion and vascular permeability which results in enhanced tissue penetration of the administered drug [18, 19].

In addition, many research data support using MHTh in combination with cancer immunotherapy. Studies have revealed its role in improved antitumor immune responses [20]. MHTh can increase the visibility of tumor cells to the immune system and enhance the trafficking of immune cells like CD8^+^ T-cell across the tumor vascular barrier to access the tumor [21, 22].

In the current study, the effect of MHTh on tumor uptake of liposomal Doxorubicin (LDox, Doxil®, excitation: 489 nm, emission: 590 nm) in a syngeneic 4T1 TNBC was investigated. Uptake of doxorubicin in control and treated tumor was investigated via *ex vivo* optical imaging of harvested tumors. Finally, we examined whether MHTh can modulate the immune microenvironment within the tumor by probing for increased transcript expression of CD8 for cytotoxic T lymphocytes (CTLs) [23], interferon-γ (IFN-γ) ^[24]^ for marking functional CTLs and PD-L1 [25] for immune checkpoint molecules that enable immune cell dysfunction.

## Materials and methods

### Cell lines and Small Animal Xenografts

All animal experiments were conducted in compliance with the animal protocol approved by the Institutional Animal Care and Use Committee (IACUC) at Wayne State University and ARRIVE 2.0 guidelines. All anesthesia and euthanasia protocols used in the study comply with the American Veterinary Medical Association (AVMA) Guidelines for the Euthanasia of Animals. 4T1 cell lines were provided by WSU Animal Model and Therapeutics Evaluation Core (AMTEC) and were grown in 5% CO_2_ using DMEM supplemented with 1% penicillin-streptomycin and 5% fetal bovine serum (Sigma) as media at 37 °C. Female BALB/c mice aged 6-8 wks (20-30 g) were purchased from Charles River Laboratories (Wilmington, MA). Mice were injected subcutaneously with 5 × 10^5^ 4T1 cells in PBS on the distal lower limb. Animals were monitored three times a week with tumors measured using a caliper and the volume determined via the formula: length × width × height × π/6. Tumor volumes ranging from 100 to 150 mm^3^ were utilized for *in vivo* treatment studies. Mice were euthanized via CO_2_ asphyxiation after intravenous injection of Hoechst 33342 as mentioned below.

### Mild Hyperthermia Treatment

A MHTh prototype thermal system was developed using a water bath sonicator (Fig. 1). The heating bath was constructed using the CREWORKS 3L ultrasonic cleaner as a base because of its size and ability to hold water at a constant 42 °C, which mimics the temperature achieved by the ultrasound hyperthermia experiments. A raised bed was constructed using metal rods, acrylic, and foam to support the mice during heating, which included mounts for anesthesia masks and holes for the insertion of the inoculated hind leg. A temperature monitoring system consisting of small footprint infrared (IR) thermometers and an Arduino UNO reading module were used to ensure the mice body temperature remained within the acceptable range in real-time. All mice in treatment (n=6) and control (n=6) groups were administered with 9 mg/kg (90 μL of 2 mg/mL) of liposomal doxorubicin hydrochloride (LDox) (Zydus, NJ, USA) via the tail vein before initiating MHTh. Immediately after the LDox injection, tumors in the treatment group were immersed in the water bath at a temperature maintained at 42 °C for 30 minutes while anesthetized with 2% isoflurane. The stage was covered with insulated material to minimize fluxes in body temperature. After MHTh was terminated, all mice received Hoechst 33342 injected through the tail vein (1 mg/100μL of saline) 5 minutes before euthanasia for nuclear staining. Tumors were harvested and fixed in formalin for further investigation.

**Figure 1.**
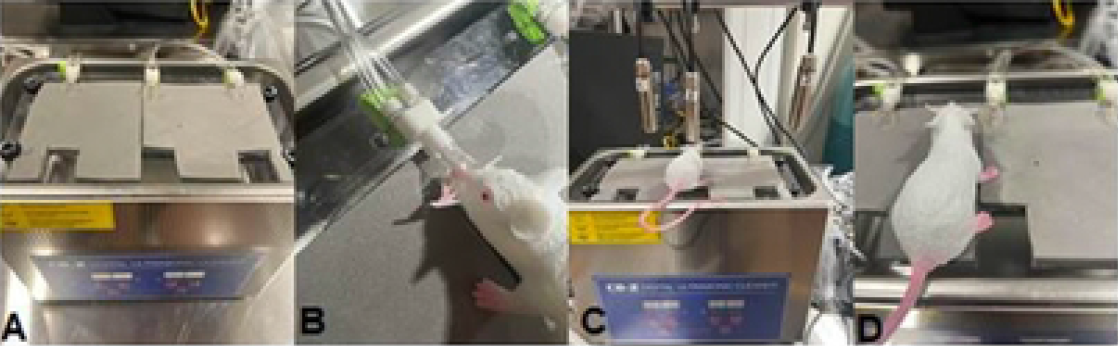
Water bath constructed unit for hyperthermia application. Tumor bearing legs were immersed in a temperature-controlled water bath (42 °C, 30 min) through an opening which enables submerging the tumor in the water bath. (A) Water bath prototype for mild hyperthermia. The gray foam serves as the mouse bed. (B) The mouse snout is placed inside the nose cone which delivers the isoflurane anesthetic. (C) Laser thermometer is positioned at the top of the animal to monitor temperature. (D) Illustration of a mouse with the hind leg inserted through the cutout where it is exposed in 42 °C.

### *Ex vivo* optical Imaging

Fluorescence images of harvested tumor from both control and MHTh treatment groups were acquired using Bruker Carestream In-Vivo Xtreme imaging system at excitation (540 nm) and emission (600 nm) wavelengths. The fluorescent intensity in each tumor was quantified using Bruker Multispectral software by manually contouring the tumor boundary.

### Fluorescent Imaging

Confocal imaging was performed on sliced tumor tissues mounted on glass slides for fluorescent visualization of LDox and Hoechst 33342. Formalin-fixed tumor tissues were replaced with ethanol (70%) before tissues were embedded in paraffin followed by sectioning. Tissues were sliced into 5 μm sections and mounted on glass slides for fluorescent microscopy (Zeiss LSM-780). Sequential sections were stained with hematoxylin and eosin (H&E) for histochemical analysis to confirm the viability of the tumor tissue. Fluorescent intensity of LDox was quantified using image J software (version 1.53K, NIH, Bethesda, USA).

### *In situ* hybridization

To evaluate immunogenic responses after MHTh treatment, we investigated the mRNA levels for CD8, IFN-γ and PD-L1 in control and treated tumor samples with RNA *in situ* hybridization (ISH) using RNAscope® kit (ACD Bio). Tumor tissue sections were subjected to single ISH detection according to manufacturer’s instruction. Positive staining was indicated by red dots present in the nucleus and/or cytoplasm. The images (40× objective) were captured under a light microscope (Carl Zeiss) and signal quantification was analyzed by QuPath software version 0.4.3.

### Statistical analysis

GraphPad Prism version 9.02 was used to perform statistical analyses unless otherwise stated. Data are presented as the mean ± S.D. Mann-Whitney t-tests were carried out for treatment and control groups studies. The significance of correlations was determined by Pearson analysis. A P value < 0.05 was considered statistically significant.

## Results and Discussion

Here, we evaluated the efficacy of MHTh on enhanced uptake of LDox, the most frequently used chemotherapy treatment for TNBC. Illustrations of a water bath unit for hyperthermia application and immersion of the target region in a temperature-controlled water bath (42 °C) is represented in Fig. 1A-D. Preparation of this prototype is straightforward with the advantage of faithfully mimicking MHTh while providing a method for isolating tumors and eliminating whole body exposure to elevated temperatures, which can be detrimental to animal health.

MHTh modulation of tumor uptake of LDox was measured and compared to the control group. *Ex vivo* imaging of harvested tumor indicates that LDox was significantly higher in MHTh-treated than untreated tumors (Fig. 2A). Quantification of fluorescent intensities of LDox in both groups indicated an enhanced uptake in mice treated with MHTh compared to the control mice that received the same dose of LDox but without any MHTh treatment (Fig. 2B) (656.5 ± 62.6 vs 770.6 ± 53.3, p<0.007).

**Figure 2.**
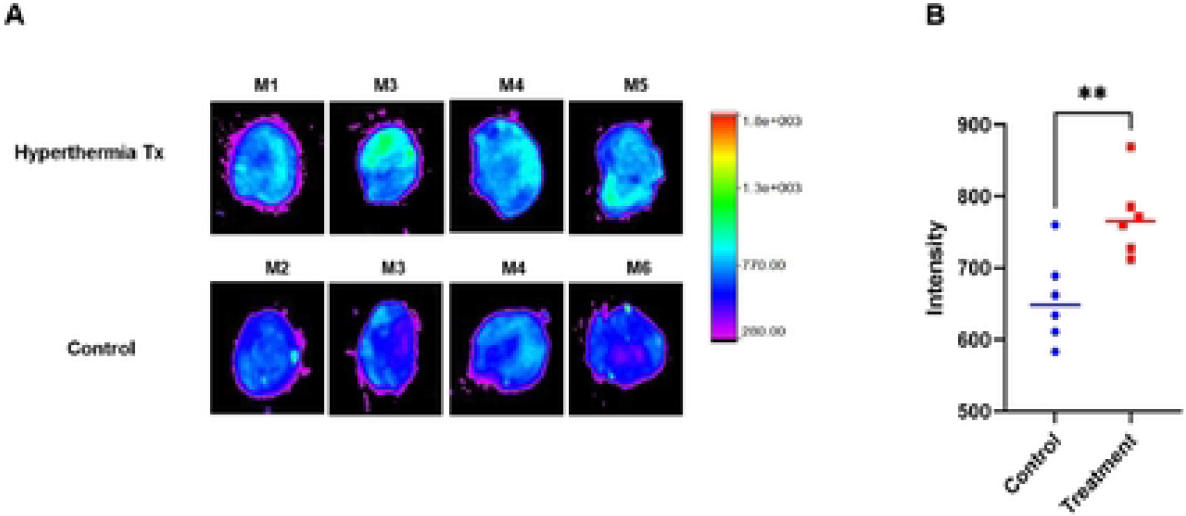
Ex vivo imaging of tumor by Carestream In Vivo Xtreme in hyperthermia treated and untreated group (A) Representative images of four different mice from treatment and control group (B) Quantified fluorescent intensity in region of interest (ROI) of tumor tissues in treatment and control group (n=6). **p < 0 01

Fluorescent microscopy of tumor tissues mounted on the slide was performed to compare uptake of Doxil in hyperthermia treatment group and control cohort. Fluorescent channels at 420-490 nm (blue) and 570-640 nm (red) were used for imaging Hoechst 33342 and Doxil respectively (Fig. 3A). Images show that in agreement with the Carestream imaging, the fluorescent intensity of red probe was significantly higher in the hyperthermia treatment group which demonstrated that hyperthermia can efficiently increase the uptake of chemotherapy agents into the tumor microenvironment. The fluorescent intensity of Doxil was quantified and data indicates higher fluorescent signal of Doxil in tumor treated compared to control (Fig. 3B). We posit that this can be due to increased blood flow, enhanced vascular permeability, and elevated metabolism level at the tumor region heated by water bath but this requires further study.

**Figure 3.**
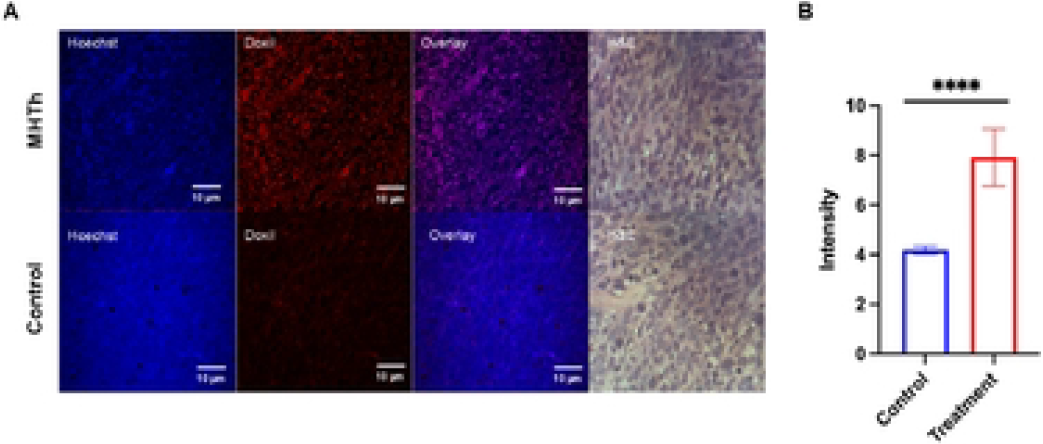
Fluorescent imaging of Doxil (red) and Hoechst 33342 (blue) in mild hyperthermia (MHTh) treatment (top) and control (bottom) groups H&E staining demonstrates tumor tissue viability. (B) Quantified fluorescent intensity of Doxil staining At least four fields per section was quantified. ****p < 0 0001

RNA ISH revealed the presence of CD8 mRNA in tumor samples treated with hyperthermia (Fig. 4). Higher transcripts (red dots) of CD8 were visible in the MHTh tumor whereas the control group had minimal to no expression. The red signal for IFN-γ were found to be very small in both control and treated groups and there was no significant difference between them. The level of PD-L1 transcript was discovered to be significantly low in the tumor sections of both the control and treated groups.

**Figure 4.**
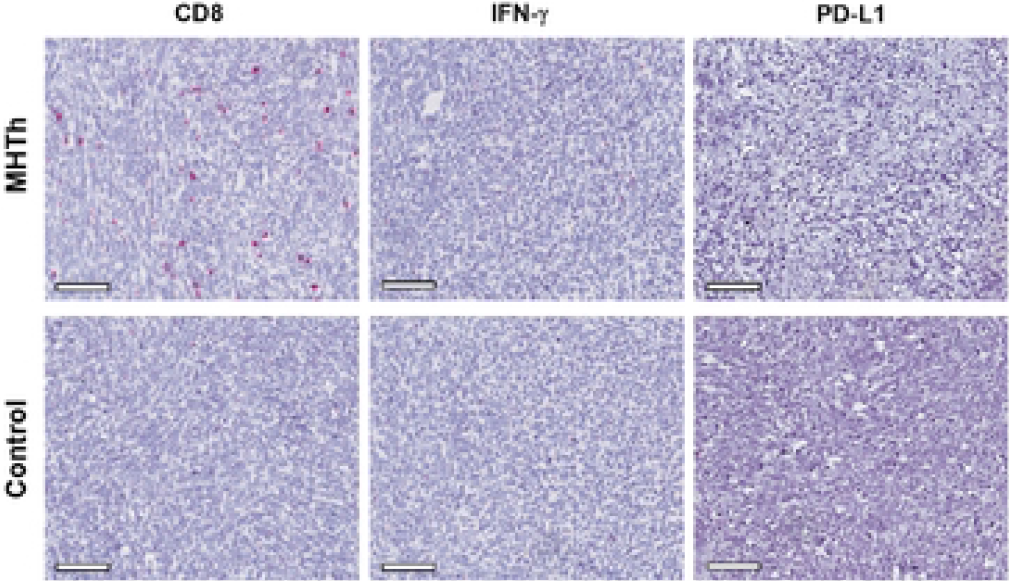
*In situ* hybridization assay to detect the expression of CD8, IFN-g and PD-L1 in control and treatment group. (A) IFN-g, CD8 and PD-L1 was detected using red chromogen. Higher transcript of CD8 was observed after hyperthermia treatment The expression of IFN-g and PD-L1 m both treated and control groups was minimal Scale bar, 100 μm.

## Conclusion

Here we showed that MHTh at 42 ºC for 30 minutes has a significant effect on the uptake of LDox as a chemotherapy agent. The application of MHTh can be beneficial for solid tumor treatment by increasing drug delivery. Moreover, we demonstrated increased infiltration of CD8 T-cells after mild hyperthermia which can be beneficial in setting of combining immunotherapy with MHTh. Our results further confirmed that MHTh to be mediated tumor infiltration of immune cells. Future studies will investigate treatment outcomes, and whether improved tumor regression is displayed by the mild hyperthermia-treated group.

## Acknowledgements

We thank the NIH NCI R37CA220482 (NTV) for funding. We also thank the Barber fellowship (WS and RS) and Wayne State University School of Medicine Summer Undergraduate Research Experience (MM). Funding was also provided by KCI Strategic Research Initiative Grant. The Microscopy, Imaging and Cytometry Resources Core (MICR) and Animal Modeling and Therapeutics Core, which provided technical assistance are supported, in part, by the NIH Cancer Center grant P30 CA022453 to KCI at Wayne State University, the Perinatology Research Branch of the National Institutes of Child Health and Development at Wayne State University.

## Author contributions

FR led the study execution, performed data analysis, wrote and revised the manuscript. WJ, RS, AP, MM, FY performed the study and assisted in writing the manuscript. AS assisted in the animal studies. MM and NTV designed and conceptualized the study. MM assisted in manuscript writing and drafting. NTV supervised the study.

## Data availability

All data required to evaluate the conclusions in the paper are present in the paper.

